# *SLC25A28* is a synthetic lethal vulnerability in nucleotide excision repair-deficient cancers

**DOI:** 10.64898/2026.10.05.756638

**Authors:** Nan Yang, Lisa Hoeg, Xuechun Bai, Sizhi Gao, Elisa de Stanchina, David Solit, Sabine Topka, Sami Belhadj, Jian Carrot-Zhang, Gopa Iyer, Daniel Durocher, Kent W. Mouw, Vijai Joseph, Kenneth Offit, Steven Lipkin

## Abstract

Approximately 10% of bladder, upper urinary tract, and endometrial cancers harbor deleterious somatic alterations in nucleotide excision repair (NER) pathway genes. Although platinum-based chemotherapy is the standard treatment for NER-deficient tumors, its utility is limited by nephrotoxicity, motivating alternative strategies that avoid direct DNA targeting. Here, we employed a genome-wide CRISPR-Cas9 screen to identify synthetic lethal vulnerabilities in NER-deficient cancer cells. An *ERCC4*-null RT112/84 bladder cancer cell line was generated and validated by Sanger sequencing and Western blotting, and functional NER deficiency was confirmed using a host cell reactivation assay. Parallel screens in *ERCC4*-deficient and isogenic wild-type cells. nominated *SLC25A28* as the top-ranked synthetic lethal candidate in *ERCC4*-deficient cells, an interaction further supported by DepMap dependency. Clonogenic assays confirmed that loss of *SLC25A28* selectively impaired cell viability in cells deficient for *ERCC2*, *ERCC3*, *ERCC4*, or *ERCC5* and inducible knockout selectively suppressed *ERCC4*-deficient xenograft growth *in-viv*o. Mechanistically, RNA sequencing followed by gene set enrichment analysis revealed enrichment of oxidative stress-related pathways, including cellular responses to reactive oxygen species (ROS) and DNA damage signatures, upon *SLC25A28* loss. These transcriptional changes are consistent with increased oxidative stress and DNA damage, placing heightened demand on NER-mediated repair. Direct measurement confirmed increased intracellular ROS. Collectively, our nominates *SLC25A28* as a previously unrecognized synthetic lethal vulnerability in NER-deficient cancers and implicate mitochondrial iron homeostasis as a candidate metabolic dependency that may be targeted independently of platinum-based chemotherapy.

**Significance:** This study identifies *SLC25A28* as a synthetic lethal vulnerability in nucleotide excision repair–deficient cancers, linking mitochondrial iron homeostasis and oxidative stress to selective DNA damage intolerance and therapeutic opportunity.

## Introduction

Nucleotide excision repair (NER) is a versatile DNA repair pathway responsible for removing bulky, helix-distorting DNA lesions induced by ultraviolet radiation(1), tobacco-derived carcinogens(2), and platinum-based chemotherapeutics(3). NER deficiency arises through inactivating alterations in core components of the pathway, including the TFIIH-associated helicases *ERCC2* (XPD) and *ERCC3* (XPB) and the structure-specific endonucleases *ERCC4* (XPF) and *ERCC5* (XPG)(4–6), and defines a distinct molecular subset of cancer(7). Large-scale genomic analyses have demonstrated that approximately 10% of bladder urothelial and endometrial cancers harbor deleterious somatic mutations in NER genes(7), establishing NER deficiency as a recurrent clinically actionable genotype in these malignancies.

In muscle-invasive urothelial carcinoma, somatic *ERCC2* mutations were among the NER alterations to be clinically validated as predictive biomarkers of cisplatin-based chemotherapy response(8–10). Platinum-based agents induce bulky DNA lesions that are exclusively and preferentially repaired by the NER pathway(3,11,12), loss-of-function *ERCC2* mutations correlate with improved pathologic complete response rates and survival in patients treated with neoadjuvant cisplatin-containing regimens(8). These observations established a clinical paradigm in which intrinsic DNA repair deficiency can be exploited therapeutically by DNA-damaging agents, while also underscoring the importance of accurately identifying NER-deficient (NER-D) tumors(8,9).

Despite this apparent platinum sensitivity, the therapeutic window for platinum-based chemotherapy in NER-D cancers remains constrained by nephrotoxicity and other systemic toxicities(13–15), particularly in older patients and those with pre-existing renal impairment(16,17). Importantly, although NER deficiency is associated with increased platinum responsiveness, the predictive utility of NER status is imperfect, and platinum resistance can arise through multiple mechanisms beyond NER deficiency(18). Thus, reliance on DNA-damaging cytotoxic agents alone provides an incomplete solution for NER-D cancers. In contrast to homologous recombination–deficient tumors, where PARP inhibition provides a clinically effective strategy that exploits DNA repair dependencies rather than relying on DNA-crosslinking cytotoxic chemotherapy(19,20), clinically actionable targeted therapies for NER-deficient tumors have not yet been established. There is therefore a critical unmet need to identify genetic vulnerabilities that selectively affect NER-D cancers while sparing normal tissues, that may be useful for further targeting.

The concept of synthetic lethality provides a powerful framework for exploiting tumor-specific DNA repair defects(21,22). Synthetic lethal interactions occur when the simultaneous disruption of two genes is lethal, whereas loss of either gene alone is tolerated, enabling selective targeting of tumor cells harboring defined genetic lesions(21,22). The clinical success of PARP inhibitors in *BRCA1*/*2* mutant tumors exemplifies the translational potential of this approach and motivates analogous efforts to identify synthetic lethal partners of NER deficiency(20,23). Complementing this genetic-dependency strategy, NER-D tumors exhibit hypersensitivity to certain genotoxic agents, including irofulven and illudin S(7,24,25), and these compounds have not yet translated into widely adopted precision therapies. Systematic identification of non-genotoxic, druggable synthetic lethal targets in NER-deficient cancers therefore remains an important unmet challenge. CRISPR–Cas9–based genome-wide loss-of-function screens enable unbiased discovery of context-specific genetic dependencies at scale(26), including synthetic lethal interactions linked to defined DNA repair defects. By comparing gene essentiality profiles between isogenic NER-proficient and NER-deficient models, such screens can uncover noncanonical vulnerabilities that arise upon loss of NER function. In interpreting candidate hits from such screens, prior evidence provides a useful framework for prioritization: oxidative stress responses can critically influence cellular tolerance to DNA damage and DNA repair defects(27–29). Within solute carrier family 25 (SLC25)(30), mitochondrial iron transporter *SLC25A28* (mitoferrin-2), mediates mitochondrial iron import(31), redox balance, and this gene family has been implicated in regulating oxidative stress–induced cell death and cancer progression. Notably, dysregulated mitochondrial iron loading can amplify DNA damage(32), suggesting a potential functional link between mitochondrial iron homeostasis, redox stress, and DNA repair capacity that could be relevant to NER-deficient cells.

In this study, we performed a genome-wide CRISPR–Cas9 screen in isogenic RT112/84 bladder cancer cells with or without *ERCC4* to systematically identify synthetic lethal partners of NER deficiency. Among the top candidates identified, *SLC25A28* emerged as a prominent hit. We therefore focused on *SLC25A28* to define the basis of its selective essentiality in NER-deficient contexts. Our data indicate that loss of *SLC25A28* selectively impairs viability across multiple NER gene knockouts, including *ERCC2, ERCC3, ERCC4, and ERCC5*. Transcriptomic analyses reveal enrichment of oxidative stress and DNA damage response signatures upon combined loss of NER function and *SLC25A28* enhances, consistent with a model in which altered mitochondrial iron handling and redox imbalance create a selective vulnerability in NER-D cells. Collectively, these findings nominate *SLC25A28* as a previously unrecognized and potentially druggable synthetic lethal vulnerability in NER-deficient cancers and suggest a prospective therapeutic strategy that decouples NER-directed precision oncology from the clinical limitations of platinum-based chemotherapy.

## Materials and Methods

### Genomic profiling and analysis of germline and somatic alterations

As part of an institutional review board–approved protocol supporting clinical management, patients with advanced cancer at Memorial Sloan Kettering Cancer Center (MSKCC) underwent matched tumor and normal DNA sequencing using the MSK-IMPACT platform, as previously described. The overall analytic workflow is summarized in Figure S1. We analyzed 95,141 patients with somatic profiling and 45,748 patients with harmonized germline reporting across 32 OncoTree tissue categories. Somatic prevalence was defined as the fraction of patients per tissue with at least one alteration in *ERCC2, ERCC3, ERCC4*, or *ERCC5*. Germline prevalence was defined as the fraction of patients per tissue with pathogenic or likely pathogenic *ERCC3* variants. We applied harmonized tissue mapping, quality-control checks for identifier consistency and duplicate handling, and tissue-level prevalence. Consent for germline testing was obtained through the referring physician after a description of the associated risks and benefits, with emphasis on patients with advanced (stage III-IV) disease, and referral remained at physician discretion. Individuals identified as carriers of pathogenic or likely pathogenic variants (hereafter “pathogenic variants”) were offered genetic counseling, and at-risk family members were identified where applicable.

For the present study, somatic variants were analyzed across all consented patients, whereas germline variants were analyzed only among the subset who consented to germline return of results. Somatic variants in *ERCC2, ERCC3, ERCC4,* and *ERCC5* were obtained from the MSKCC cBioPortal, and their prevalence was determined separately for each gene across tumor types. Germline analysis was restricted to *ERCC3*, the only NER gene included on the germline panel; germline *ERCC3* variants were interpreted according to American College of Medical Genetics and Genomics (ACMG) criteria by a clinical molecular geneticist or molecular pathologist, and variants of uncertain significance were not included in clinical reporting.

Because a subset of patients had more than one tumor sequenced, records were merged and deduplicated so that each patient contributed only once to the somatic denominator. For patients with multiple sequenced tumors, the tumor harboring an *ERCC2*/3/4/5 alteration was preferentially retained to avoid undercounting carriers; germline variants were counted once. Following deduplication, variants were restricted to the *ERCC* genes of interest and stratified by cancer type, and prevalence was computed separately for somatic *ERCC2/3/4/5* and germline *ERCC3* (Figure 2). Germline and somatic prevalence were therefore calculated against distinct denominators: germline among the ∼12,000 patients consented to germline testing, and somatic among all ∼78,000 patients consented to the primary tumor: normal sequencing protocol.

### Generation of KU1919 *ERCC5* knockout cell lines

Guide RNAs targeting *ERCC5* were cloned into the px330-mCherry plasmid (Addgene, Cat# 98750). The resulting plasmids were transfected into KU1919 wild-type cells using Lipofectamine™ 3000 according to the manufacturer’s instructions. Transfected cells were sorted based on mCherry fluorescence and seeded as single cells into 96-well plates. Single-cell–derived colonies were expanded and screened by genotyping using Sanger sequencing. Guide RNA sequences are listed in Supplementary Table S1.

### Sanger sequencing

The Cas9-edited genomic regions were amplified by PCR, and PCR products were analyzed by Sanger sequencing. Primers used for gene amplification and sequencing are listed in Supplementary Table S2.

### Western blot

Cells were lysed using RIPA Lysis Buffer (Thermo Fisher Scientific, Cat# 89901) supplemented with protease inhibitors (Thermo Fisher Scientific, Cat# 78429) according to the manufacturer’s instructions. Protein concentrations were determined using the DC Protein Assay (Bio-Rad, Cat# 5000116) according to the manufacturer’s instructions. Postnuclear supernatants were obtained by centrifugation at 15,700 × g for 15 min at 4 °C and were then mixed with SDS sample buffer (Thermo Fisher Scientific, Cat# NP0007) and heated at 70 °C for 10 min. Proteins were resolved on NuPAGE 4%–12% Bis-Tris gels (Thermo Fisher Scientific) and transferred to PVDF membranes. Primary and secondary antibodies used are listed in Supplementary Table S3.

### Host cell reactivation assay

DNA nucleotide excision repair (NER) capacity in cells was assessed using a host cell reactivation assay adapted from previously published protocols(33). 3 × 10^5^ cells were seeded in 6-well plates and transfected the next day with BFP (800J/m^2^ UV treatment) and mcherry plasmids (internal control) using Lipofectamine 3000 (Thermo Fisher Scientific, Cat# L3000015). each transfection was performed in triple replication. Twenty-four hours post-transfection, cells were harvested, resuspended in PBS containing 2% FBS and 5 mM EDTA, optionally stained with To-Pro-3 (Thermo Fisher Scientific, Cat T3605) filtered, and analyzed by flow cytometry. Quantitation:

Expression of the UV-treated BFP reporter in cells transfected with the damaged plasmid cocktail was normalized to the expression of the co-transfected undamaged mcherry reporter, calculated as: X= (BFP Count* Mean BFP intensity)/(mcherry Count* Mean mcherry intensity).Expression of the unmodified BFP reporter in cells transfected with the undamaged plasmid cocktail was similarly normalized to the co-transfected undamaged mcherry reporter: Y=(BFP Count* Mean BFP intensity)/(mcherry Count* Mean mcherry intensity). The relative reporter expression was calculated as the ratio of normalized BFP expression in the damaged condition (X) to that in the undamaged condition (Y), multiplied by 100 to obtain the percentage of reporter expression: Z =100*X/Y.

### Genome-wide CRISPR–Cas9 knockout screen and data analysis

Genome-wide CRISPR–Cas9 loss-of-function screening was performed following established approaches with modifications as described below(26). RT112/84 wild-type and *ERCC4*-knockout isogenic cell lines were first transduced with lentiCas9-Blast (Addgene, #52962) to generate stable cell populations with constitutive Cas9 expression. Cas9-expressing cells were subsequently infected with the human Brunello sgRNA library at a multiplicity of infection (MOI) of 0.3 to ensure predominantly single sgRNA integration per cell. Transduced cells were subjected to puromycin selection (1 μg/mL) for 3 days to enrich for sgRNA-positive cells. Cells harvested at day 3 post-infection were designated as the baseline reference population (T0). At this time point, cells were pooled and evenly split into three technical replicates for each genotype. Cells were then propagated for an additional 18 days under conditions maintaining a minimum representation of 1,000 cells per sgRNA throughout the screen. At the endpoint, cell pellets were collected from each replicate for genomic DNA extraction. Integrated sgRNA cassettes were PCR-amplified from genomic DNA and prepared for next-generation sequencing.

Read counts were generated from fastq files using MAGeCK(34). The mean of T0 replicates was used as the control for subsequent analyses. BAGEL2, a Bayesian Analysis of Gene Essentiality(35), and Crispr Counts Analysis (CCA)(36), were used for testing essentiality and fitness differences of genes in the experimental conditions.

### Publicly available cancer gene essentiality datasets

Gene-level fitness effects across diverse cancer cell lines were assessed using large-scale CRISPR screening data obtained from publicly accessible resources. Chronos dependency scores were retrieved from the 2025 Q3 release of the Broad Cancer Dependency Map and used to quantify gene essentiality across the cell line panel.

### Lentiviral transduction

Lentiviral particles were generated in 293T cells cultured in 10 cm dishes. Cells were co-transfected with psPAX2, pMD2.G, and lentiCRISPR v2 plasmids (Addgene, Cat# 12260, #12259, #52961) encoding the desired sgRNA using X-tremeGENE transfection reagent (Roche, Cat# 6366236001). Virus-containing supernatants were collected at 48 and 72 hours post-transfection. Target cells were then infected in the presence of polybrene (Sigma-Aldrich, Cat# TR-1003) to enhance viral entry.

### Colony formation assay

Cells were seeded at the specified density and transduced overnight with lentivirus encoding sgRNAs targeting genes of interest in the presence of 5–7.5 μg/mL polybrene (Sigma-Aldrich, Cat# TR-1003). At least two independent sgRNAs were used to validate each target gene. Approximately 14 h post-transduction, the medium was replaced, and cells were selected using 1 μg/mL puromycin for 3 days. After selection, cells were detached using 0.25% trypsin, neutralized with medium containing 10% FBS, and live cells were counted. Cells were then plated at low density in 6-well or 10 cm plates in triplicate and cultured until colonies contained more than 50 cells. Colonies were fixed and stained with a solution of 6% glutaraldehyde and 0.5% crystal violet for at least 30 min, gently rinsed with tap water, and air-dried at room temperature. Colonies were counted manually under a stereomicroscope, and surviving fractions were calculated relative to the plating efficiency of untreated control cells.

### RNA isolation, RNA sequencing (RNA-seq), and bioinformatics analysis

RNA purification, library preparation, and sequencing were performed by the Memorial Sloan Kettering Cancer Center Integrated Genomics Operation (IGO) Core. Total RNA was isolated using a combined TRIzol and column-based purification approach. Briefly, cells were lysed in 1 mL TRIzol Reagent (Thermo Fisher Scientific, catalog #15596018), and phase separation was induced by addition of 200 µL chloroform. The aqueous phase was collected, and RNA was purified using the miRNeasy Mini Kit (Qiagen, catalog #217004) on a QIAcube Connect (Qiagen) according to the manufacturer’s protocol. RNA was eluted in 54 µL RNase-free water.

RNA concentration was determined using the RiboGreen assay, and RNA integrity was assessed using an Agilent BioAnalyzer. A total of 500 ng of high-quality RNA (RIN >9.5) was subjected to poly(A) selection and library preparation using the Watchmaker mRNA Library Prep Kit (catalog #7BK0001) according to the manufacturer’s instructions, with 8 cycles of PCR amplification. Libraries were barcoded and sequenced on an Illumina NovaSeq X platform using paired-end 100-bp reads (NovaSeq X 10B Reagent Kit). An average of 94 million paired-end reads was generated per sample. Ribosomal RNA reads accounted for 0.10–0.13% of total reads, and the percentage of bases mapping to mRNA averaged 89%.

RNA-seq data analysis was performed by the MSKCC Bioinformatics Core using standardized workflows. Raw sequencing reads (FASTQ files) were aligned to the human reference genome (UCSC hg38) using STAR, and gene-level expression counts were generated from the aligned reads. Count data were normalized and differential gene expression analysis was performed using DESeq2. Differentially expressed genes were identified based on adjusted P values (FDR < 0.05) and fold-change thresholds as indicated. Pathway enrichment analysis was conducted using Gene Set Enrichment Analysis (GSEA), with genes ranked by differential expression statistics and evaluated against curated gene sets from the Molecular Signatures Database (MSigDB).

### Assessment of Cellular Reactive Oxygen Species (ROS)

To assess reactive oxygen species (ROS) levels, cells were incubated with 10 μM 2′,7′-dichlorofluorescin diacetate (DCFDA) in culture medium for 30 minutes at 37 °C, followed by washing with PBS. Cells were then trypsinized, harvested, and centrifuged to remove trypsin, before being resuspended in staining buffer containing 50 nM To-Pro-3 (Thermo Fisher Scientific, Cat# T3605). The cell suspensions were filtered through a 40 μm cell strainer, and were analyzed by flow cytometry.

### Animals

All animal experiments were conducted in accordance with protocols approved by the Institutional Animal Care and Use Committee of Memorial Sloan Kettering Cancer Center (MSKCC). Mice were housed in a temperature-and humidity-controlled animal facility under a 12-hour light/12-hour dark cycle. Animals had ad libitum access to standard laboratory chow (LabDiet 5053) and water throughout the study. Mice were group housed in individually ventilated cages with autoclaved corncob bedding and allowed to acclimate to the facility for a minimum of 5 days prior to experimental procedures.

### Xenograft studies

A doxycycline-inducible CRISPR–Cas9 system (TLCV2, Addgene #87360) was used to generate lentivirus encoding an sgRNA targeting *SLC25A28* or a non-targeting control sgRNA. RT112/84 wild-type (WT) and RT112/84 *ERCC4*-knockout cells were transduced with the corresponding lentiviruses. Approximately 14 h after infection, the medium was replaced, and transduced cells were selected with puromycin (1 μg/mL) for 3 days. Following selection, cells were expanded in culture and used for subsequent xenograft experiments.

Female NSG mice (6–8 weeks old; The Jackson Laboratory) were maintained under institutionally approved conditions for in vivo studies. Subcutaneous xenografts were established by injecting 5×10^6^ cells suspended in Matrigel into both flanks of each mouse (n = 6 mice per group). Tumor growth was monitored, and once tumors reached an average volume of approximately 50–200 mm³, mice were placed on a doxycycline-containing diet (Inotiv, Cat# D18042704i). Tumor dimensions were measured twice weekly using calipers, and tumor volume was calculated as length × width² × 0.52. Body weight was recorded concurrently to monitor animal health. Mice were euthanized when tumor volume reached approximately 2,000 mm³.

### T7 endonuclease I (T7EI) mismatch-cleavage assay

Genomic DNA was extracted from CRISPR-edited cells. Target genomic loci flanking the sgRNA cut site were amplified by PCR. PCR primers were designed to generate amplicons, with the Cas9 cleavage site placed off-center within the amplicon T7EI-based editing-efficiency detection was performed using the Alt-R™ Genome Editing Detection Kit (Integrated DNA Technologies, Cat #1075931) following the manufacturer’s protocol. Briefly, PCR products were denatured and slowly re-annealed to form wild-type/mutant heteroduplexes, then digested with T7EI endonuclease I. Digested fragments were separated by agarose-gel electrophoresis. PCR primer sequences are listed in Supplementary Table S2.

### Statistical analysis

The number of experimental replications and mice used are described in figure legends. Statistical analyses were performed using GraphPad Prism software (version 8). Differences in HCRA between RT112/84 WT and RT112/84 *ERCC4*^−/−^ cell lines were assessed using a two-tailed Student’s t-test. Comparisons of dependency scores between *ERCC4* wild-type and *ERCC4* mutant cell lines in the Broad DepMap datasets were performed using the Mann–Whitney U test. For colony formation assays, comparisons between two groups were conducted using a Student’s t-test with Welch’s correction. A P value < 0.05 was considered statistically significant.

## Results

### Prevalence and spectrum of functionally disruptive NER pathway alterations in patients with cancer

Nucleotide excision repair (NER) deficiency represents a clinically important targetable vulnerability, particularly for synthetic lethal therapeutic strategies. Landmark studies using the MSK-IMPACT platform have established the prevalence and spectrum of functionally disruptive NER pathway alterations across human cancers, providing critical proof of principle that NER defects are common in malignant disease(7). However, given the extensive molecular and clinical heterogeneity of cancer, further validation and refinement in a substantially larger patient population are needed to generate more precise frequency estimates, enhance resolution across diverse histologies, and strengthen the translational rationale for NER-directed synthetic lethal approaches. Somatic *ERCC2/3/4/5* alterations were distributed across a broad range of cancer types, with the highest total prevalence in non-melanoma skin cancer (∼18%) and bladder cancer (∼17%) (Fig. 1A), followed by endometrial (14%), small bowel (12%) and melanoma (10%). The gene composition of these alterations was strongly tumor-type-dependent: bladder cancer was distinguished by a high frequency of *ERCC2* alterations (∼11% of patients), consistent with the established role of *ERCC2* in this urothelial carcinogenesis, whereas non-melanoma skin cancer was dominated by *ERCC4* and *ERCC5* alterations (5% each). Endometrial, small bowel, and colorectal cancers also showed appreciable cumulative ERCC-alteration prevalence and relatively balanced contributions across the four genes. These data confirm that somatic NER-gene alterations define a recurrent molecular subset across multiple cancer types, with bladder cancer showing the highest *ERCC2*-specific prevalence, providing the clinical rationale for identifying selective vulnerabilities in NER-deficient tumors.

**Figure 1.**
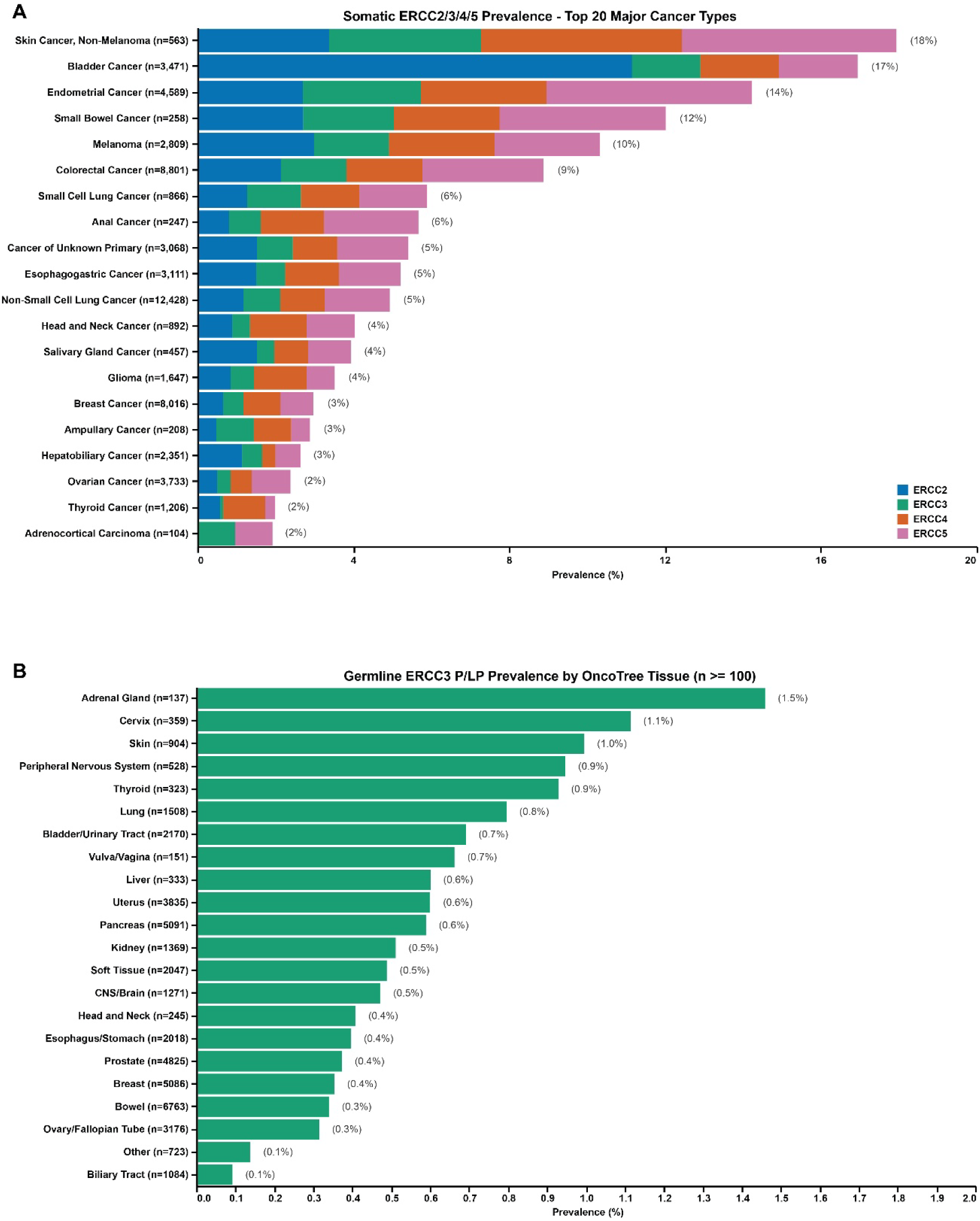
Prevalence and distribution of functionally relevant alterations in NER pathway components *ERCC2–5* among human cancers. (A) Pan-cancer somatic alteration frequency of functionally impactful variants in ERCC2–5 across tumor types annotated using the MSK-OncoTree tumor classification ontology, in a cohort of 95,141 patients profiled by MSK-IMPACT. (B) Landscape of germline alterations with predicted functional impairment in *ERCC3* variants within the same patient population.

In this expanded cohort, among NER genes with clinical reporting on the MSK-IMPACT assay, germline pathogenic/likely pathogenic NER alterations were identified in *ERCC3* in adrenal gland tumors (1.5%), cervical cancer (1.1%) and skin cancer (1.0%) (Fig. 1B). Collectively, these data confirm and refine the landscape of NER deficiency across human cancers, yielding more robust and precise prevalence estimates than previously reported from 94,098 patients spanning 32 major cancer tissue categories as defined by the OncoTree classification system(37). These observations solidify the clinical relevance of NER pathway inactivation and provide a rigorous, large-scale foundation for investigating synthetic lethal therapeutic strategies in NER-deficient tumors. Strikingly, among 217 patients carrying germline pathogenic/likely pathogenic ERCC3 variants, only 4 (1.84%) exhibited a second somatic hit in the same gene. This low frequency of biallelic inactivation is markedly inconsistent with the Knudson two-hit hypothesis for classical tumor suppressors(38). This paucity of biallelic inactivation, despite a large cohort and deep sequencing coverage, suggests that heterozygous NER dysfunction may be sufficient to confer a tumorigenic phenotype, consistent with a dominant-negative mechanism in which mutant ERCC3/XPB subunits poison the TFIIH complex and impair NER activity without requiring loss of heterozygosity(39,40). Although *ERCC2* is the predominant NER alteration in bladder cancer, we used a cleaner *ERCC4*-null model as a tractable, complete-loss-of-function system, and subsequently extended validation across *ERCC2/3/5* to test synthetic lethality against NER deficiency broadly.

### Cell lines and generation of *ERCC4^−/−^*

*ERCC4* was knocked out in RT112/84 wild-type (WT) cells using CRISPR–Cas9, generating isogenic NER-deficient cells. Sanger sequencing confirmed a homozygous frameshift mutation in exon 1 (c.27_48delGATTGCCATGGCGCCGCTGCTG, p.I10S*16) (Fig. 2A), and Western blot showed complete loss of ERCC4 protein (Fig. 2B). Functional assessment by host cell reactivation assay (HCRA) demonstrated markedly reduced NER activity in RT112/84 *ERCC4^−/−^*cells compared with WT (5.5% vs 32.9%) (Fig. 2D), confirming that RT112/84 *ERCC4^−/−^* cells are NER-deficient. An independent *ERCC4^−/−^* model generated in the H460 background was similarly validated by Sanger sequencing, Western blotting, and HCRA (Fig. S2C–E), mitigating potential clonal artifacts.

**Figure 2.**
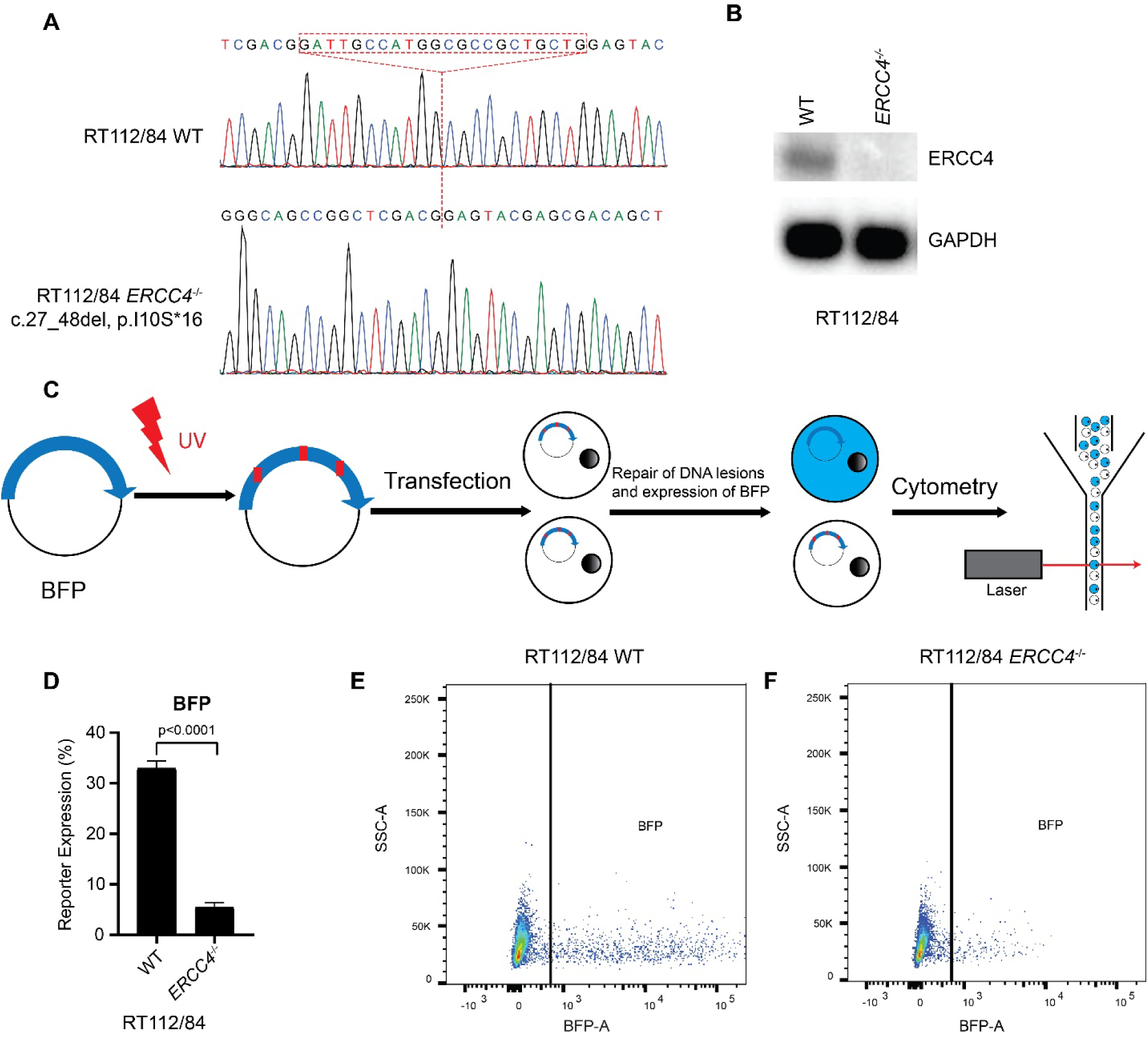
Generation of RT112/84 WT and RT112/84 *ERCC4^−/−^* isogenic cell lines and assessment of NER DNA repair capacity using the Host Cell Reactivation Assay (HCRA). (A) Sanger sequencing confirms the *ERCC4* frameshift mutation c.27_48delGATTGCCATGGCGCCGCTGCTG, p.I10S*16 in exon 1 of the RT112/84 *ERCC4^−/−^* cell line. (B) Western blot analysis shows loss of ERCC4 protein expression in RT112/84 *ERCC4^−/−^* cells. (C) Schematic workflow of the HCRA used to assess NER repair activity. (D) HCRA results demonstrate that the NER repair capacity of RT112/84 *ERCC4^−/−^* cells is significantly lower than that of RT112/84 WT cells. (E, F) Representative flow-cytometry plots from the Host Cell Reactivation Assay.

### CRISPR screen identifies *SLC25A28* as synthetic lethal with *ERCC4*

To identify genes essential for the viability of *ERCC4*-deficient cells, we performed genome-scale dropout CRISPR-based synthetic lethal screens in isogenic RT112/84 wild-type (WT) and *ERCC4^−/−^*cells. Screens were performed with the human Brunello sgRNA library and analyzed via a customized pipeline, BAGEL2(35), and CRISPR count analysis (CCA)(36) to detect genes selectively essential in the mutant cell line. Synthetic-lethal hits were defined by a CCA score >1.0 and Bayes factor (BF) >10. This filtering identified 8 genes whose loss selectively reduced fitness in ERCC4-deficient cell (Fig. 3B).

**Figure 3.**
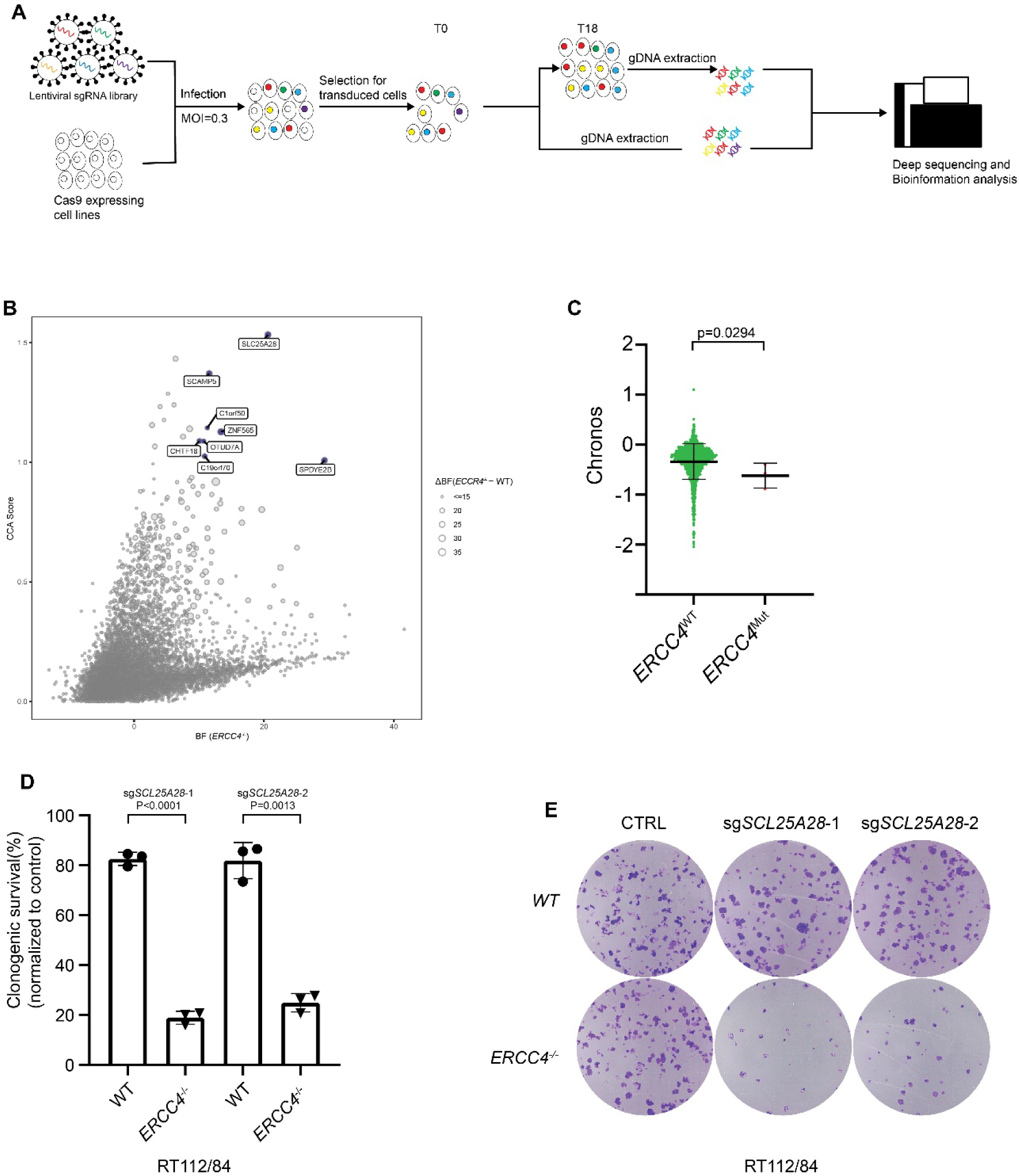
Identification of a synthetic-lethal interaction between *SLC25A28* and *ERCC4*. (A) Schematic of the whole-genome CRISPR–Cas9 screen performed in RT112/84 WT and RT112/84 *ERCC4^−/−^* isogenic cell lines. (B) Scatter plot showing CRISPR count analysis (CCA) scores and Bayes factor (BF) values generated by BAGEL2 for RT112/84 isogenic cell lines. (C) Analysis of Broad DepMap datasets comparing dependency scores between *ERCC4* wild-type (n = 1183) and *ERCC4* mutant (n = 3) cell lines. Statistical significance was assessed using pairwise Mann–Whitney U-tests. (D) Colony-formation assays of RT112/84 WT and RT112/84 *ERCC4^−/−^* cell lines transduced with the indicated sgRNAs or with sgNon-targeting (CTRL) lentivirus. Bars represent mean ± s.d. (n = 3). (E) Representative images of colony-formation assays corresponding to (D).

To prioritize genetic interactions most relevant to the tumor context, we next analyzed large-scale cancer dependency data from the DepMap projects(41,42). Cancer cell lines were stratified based on the presence or absence of damaging *ERCC4* alterations (n = 3 mutant vs. 1,183 WT), and gene-level depletion scores were compared between groups, with lower scores indicating greater negative effects on cell fitness. Although the dataset contained only a limited number of *ERCC4*-mutant cell lines, depletion of *SLC25A28* resulted in the most pronounced reduction in fitness specifically in *ERCC4*-deficient cells (Fig. 3C).

Based on these findings, we performed clonogenic survival assays to validate the synthetic lethality between *SLC25A28* and *ERCC4* in engineered RT112/84 *ERCC4^−/−^* cells. Loss of *SLC25A28*, achieved using a highly efficient sgRNA that robustly depleted *SLC25A28* expression (Fig. S3A–D), markedly reduced clonogenic survival in *ERCC4*-deficient cells compared with WT controls (Fig. 3D–E). To assess whether this interaction was cell line–specific, we extended our analysis to an additional isogenic pair, H460 WT and H460 *ERCC4*^−/−^ cells, in which *ERCC4* knockout was confirmed by Sanger sequencing and Western blotting and shown to confer functional NER deficiency by host cell reactivation assay (HCRA) (Fig. S2C-E). Consistently, *SLC25A28* loss selectively impaired the viability of *ERCC4*-deficient H460 cells (Fig. S2A, B), further confirming a synthetic lethal interaction between *SLC25A28* and *ERCC4*.

To determine whether the synthetic lethal interaction between *SLC25A28* and *ERCC4* observed in vitro is maintained in a physiological tumor context, we evaluated this interaction in vivo using a xenograft model. A doxycycline-inducible CRISPR–Cas9 system was used to generate lentivirus encoding an sgRNA targeting *SLC25A28* or a non-targeting control. RT112/84 WT and RT112/84 *ERCC4*-knockout cells were transduced and implanted for xenograft studies. In the RT112/84 WT background, inducible knockout of *SLC25A28* did not significantly affect tumor growth compared with the non-targeting control (Fig. 6B). In contrast, inducible knockout of *SLC25A28* in RT112/84 *ERCC4*-knockout tumors resulted in a significant inhibition of tumor growth relative to control tumors (Fig. 6C). Collectively, these in vivo results further support that *SLC25A28* and *ERCC4* constitute a synthetic lethal gene pair.

### CRISPR screen identifies *SLC25A28* as synthetic lethal with NER-D

To determine whether the synthetic lethal interaction extends beyond *ERCC4* to NER deficiency more broadly, we generated *ERCC2*-, *ERCC3*-, and *ERCC5*-mutant cell lines. *ERCC2* (XPD) and *ERCC3* (XPB) are essential TFIIH components required for transcription and nucleotide excision repair and show strong dependency across cancer cell lines in DepMap analyses. Consistent with this essentiality, repeated CRISPR–Cas9 targeting did not yield viable clones with biallelic disruption, and only heterozygous knockout cell lines could be established. Accordingly, RT112/84 *ERCC2^+/–^* and RT112/84 *ERCC3^+/–^*cell lines were generated in the RT112/84 background using CRISPR–Cas9. In contrast, *ERCC5* is dispensable for cell viability, permitting the generation of *ERCC5^−/−^* cells in the KU1919 background.

Heterozygous edits of *ERCC2* were confirmed by Sanger sequencing (Fig. S4A), and Western blot analysis demonstrated reduced *ERCC2* protein expression in RT112/84 *ERCC2^+/–^* cells compared with WT controls (Fig. 4C). Similarly, heterozygous knockout of *ERCC3* was validated by Sanger sequencing (Fig. S4B–C), with correspondingly decreased ERCC3 protein levels in RT112/84 *ERCC3^+/–^*cells relative to WT cells (Fig. 4D). Complete knockout of *ERCC5* was confirmed by Sanger sequencing in KU1919 *ERCC5^−/−^*cells (Fig. S4D).

**Figure 4.**
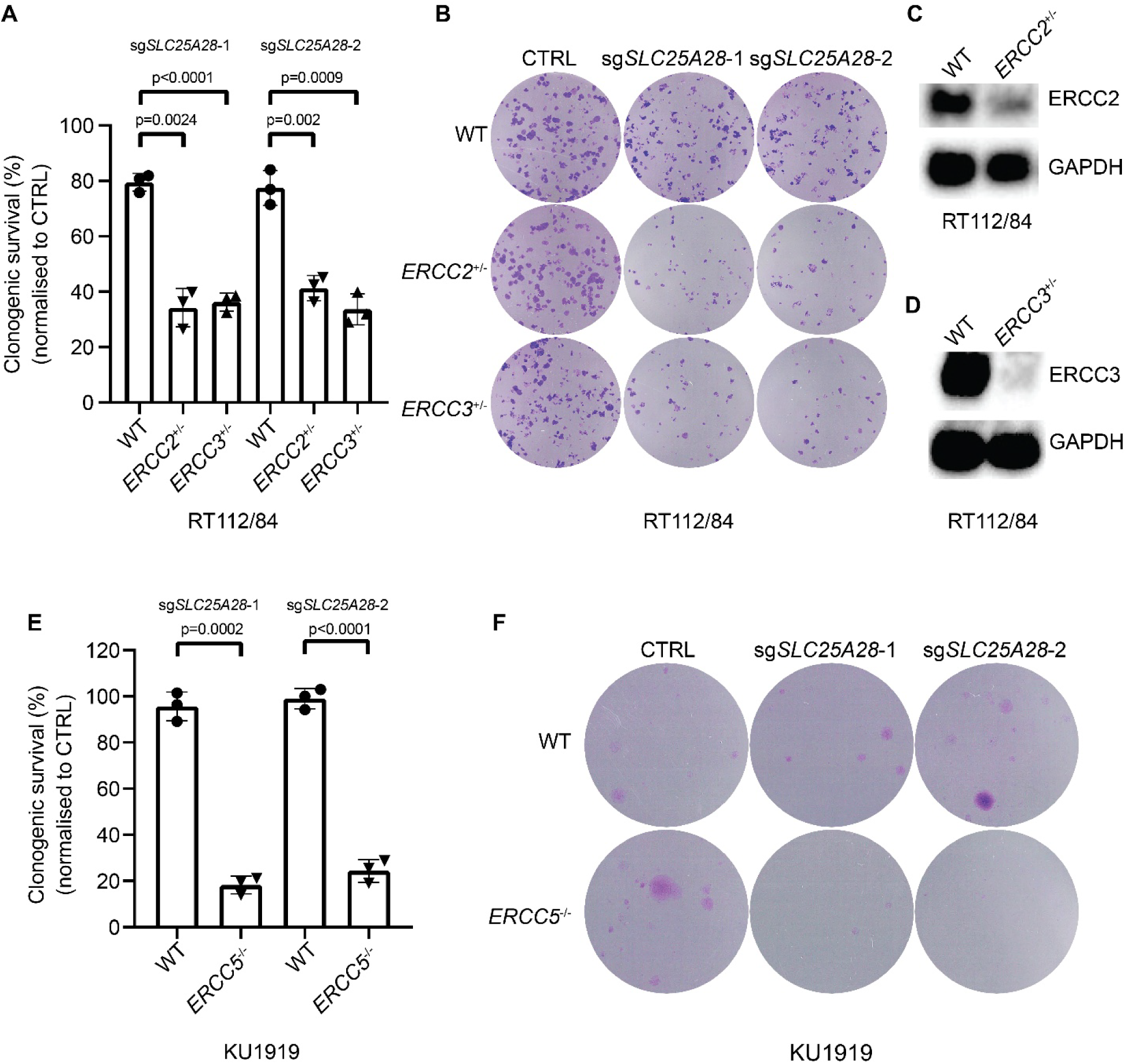
Identification of synthetic-lethal interactions between *SLC25A28* and *ERCC2*, *ERCC3*, and *ERCC5*. (A) Colony-formation assays of RT112/84 WT, RT112/84 *ERCC2^+/−^*and RT112/84 *ERCC3^+/−^* cell lines transduced with the indicated sgRNAs or with sgNon-targeting (CTRL) lentivirus. Bars represent mean ± s.d. (n = 3). (B) Representative images of colony-formation assays corresponding to (A). (C,D) Western blot analysis confirming loss of ERCC2 and ERCC3 protein expression in RT112/84 *ERCC2^+/–^*and RT112/84 *ERCC3^+/–^* cells, respectively. (E) Colony-formation assays of KU1919 WT, KU1919 *ERCC5^−/−^* cell lines transduced with the indicated sgRNAs or with sgNon-targeting (CTRL) lentivirus. Bars represent mean ± s.d. (n = 3). (F) Representative images of colony-formation assays corresponding to (E).

Clonogenic survival assays demonstrated that loss of *SLC25A28* markedly reduced clonogenic survival in RT112/84 *ERCC2^+/–^* cells compared with WT controls (Fig. 4A–B). A similar reduction in clonogenic survival was observed in RT112/84 *ERCC3^+/–^* cells following *SLC25A28* loss (Fig. 4A–B). In addition, loss of *SLC25A28* significantly impaired clonogenic survival in KU1919 *ERCC5^−/−^* cells compared with KU1919 WT cells (Fig. 4E-F). Germline loss-of-function variants in *ERCC3* have been previously shown to confer functional nucleotide excision repair deficiency, including hypersensitivity to the transcription-coupled repair-specific genotoxins illudin-S and irofulven(7,25). Helicase-domain ERCC2 alterations similarly confer functional NER deficiency(6). The reduced-*ERCC3*/4 lines generated here therefore recapitulates a genotype with an established NER-deficient functional consequence. Collectively, these results support a synthetic lethal interaction between *SLC25A28* and NER deficiency.

### *SLC25A28* deficiency increases reactive oxygen species and induces DNA damage

To investigate the mechanism underlying the synthetic lethal interaction between *SLC25A28* loss and NER deficiency, we performed RNA sequencing (RNA-seq) analysis. RT112/84 *ERCC4^−/−^* cells were transduced with a lentiviral sgRNA targeting *SLC25A28*, while cells transduced with a non-targeting sgRNA served as controls. Gene set enrichment analysis (GSEA) revealed significant enrichment of pathways related to oxidative stress, including cellular response to hydrogen peroxide, reactive oxygen species, and oxidative stress–associated gene signatures (Fig. 5A–D). In addition, the Chuang oxidative stress response pathway was significantly enriched (Fig. 5A). These transcriptional changes are consistent with increased intracellular ROS levels following *SLC25A28* loss. GSEA also showed enrichment of DNA damage–associated gene sets, including the Kyng DNA damage-downregulated signature (Fig. 5C), as well as UV response pathways such as the Gentile UV high-dose downregulated and hallmark UV response up gene sets (Fig. 5D). Given that NER is the primary pathway responsible for the repair of UV-induced DNA lesions, enrichment of these pathways suggests increased DNA damage burden and engagement of the NER pathway, though DNA damage was not directly measured. Collectively, these RNA-seq data indicate that loss of *SLC25A28* leads to elevated oxidative stress, resulting in increased DNA damage that requires NER-mediated repair.

**Figure 5.**
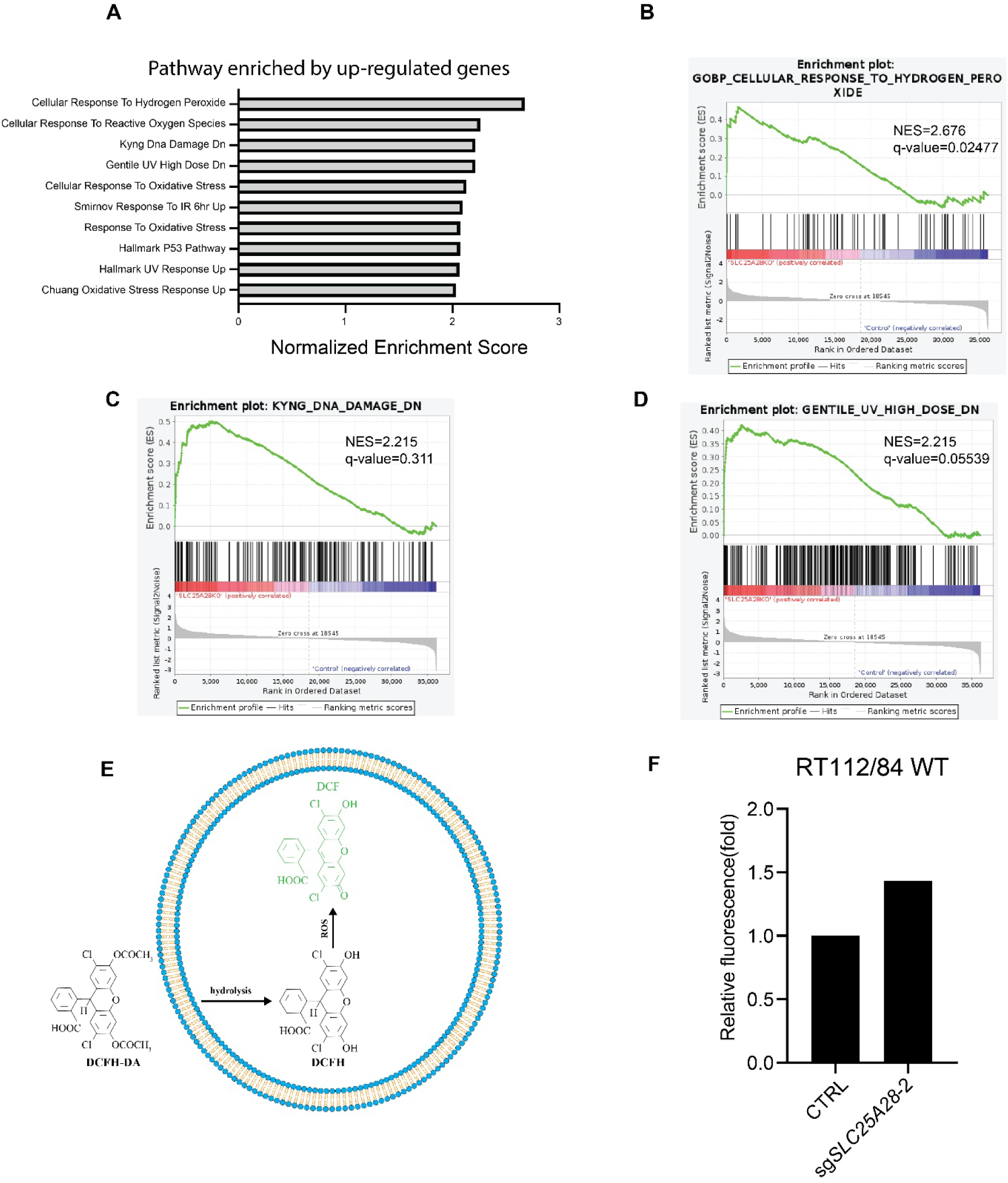
*SLC25A28* deficiency increases reactive oxygen species and induces DNA damage. (A) GSEA identified significantly upregulated pathways (FDR < 0.05). Absolute normalized enrichment scores from the comparison of *SLC25A28* KO vs. Ctrl in RT112/84 *ERCC4^−/−^* cells are shown. (B–D) Gene Set Enrichment Analysis (GSEA) of bulk RNA-seq data from *SLC25A28* KO vs. Ctrl. (E) Schematic of the DCFDA assay for measuring intracellular ROS. (F) Quantification of ROS levels after *SLC25A28* knockout in RT112/84 WT cells.

To directly validate ROS induction, *SLC25A28* was knocked out in RT112/84 wild-type cells, and intracellular ROS levels were measured using the DCFDA assay. Consistent with the transcriptomic analysis, *SLC25A28* knockout resulted in a significant increase in ROS levels compared with control cells (Fig. 5E-F).

## Discussion

In this study, we identified *SLC25A28* as a novel candidate synthetic lethal vulnerability in NER-deficient cancers. Using colony formation assays, we demonstrated that loss of *SLC25A28* reduced clonogenic survival of cells with reduced or absent functions of *ERCC2*, *ERCC3*, *ERCC4*, or *ERCC5*, consistent with a synthetic lethal interaction with NER deficiency. Consistently, *in vivo* xenograft studies revealed that inducible knockout of *SLC25A28* significantly inhibited tumor growth in RT112/84 *ERCC4*^−/−^ tumor, but did not significantly affect *ERCC4*-proficient tumors. Our Mechanistic data are consistent with a model in which loss of *SLC25A28* perturbs mitochondrial redox balance, which we propose imposes and increased DNA damage burden, converting endogenous oxidative stress into cytotoxic DNA damage (Fig. 6D). The resulting selective lethality in NER-deficient cell, contrasts to conventional chemotherapy which induces exogenous DNA damage. Somatic nucleotide excision repair deficiency is prevalent among bladder and upper urinary tract cancers, as well as endometrial cancers(7), and is associated with enhanced sensitivity to platinum-based chemotherapy(8). However, the clinical utility of platinum is constrained by nephrotoxicity, particularly in older patients or those with compromised renal function(13,16,17). The synthetic lethality paradigm, exemplified by PARP inhibitors in BRCA1/2-deficient tumors, demonstrates the potential of exploiting DNA repair deficiencies therapeutically(19). Previous NER-targeted approaches, including irofulven and illudin S, have shown preclinical efficacy but have not translated into widely adopted precision therapies(7,43,44). Among other candidates identified in our genome-wide CRISPR screen, *SLC25A28* exhibited the most consistent synthetic lethal effect, supporting its prioritization for further investigation as a candidate vulnerability in NER-deficient cancers.

**Figure 6.**
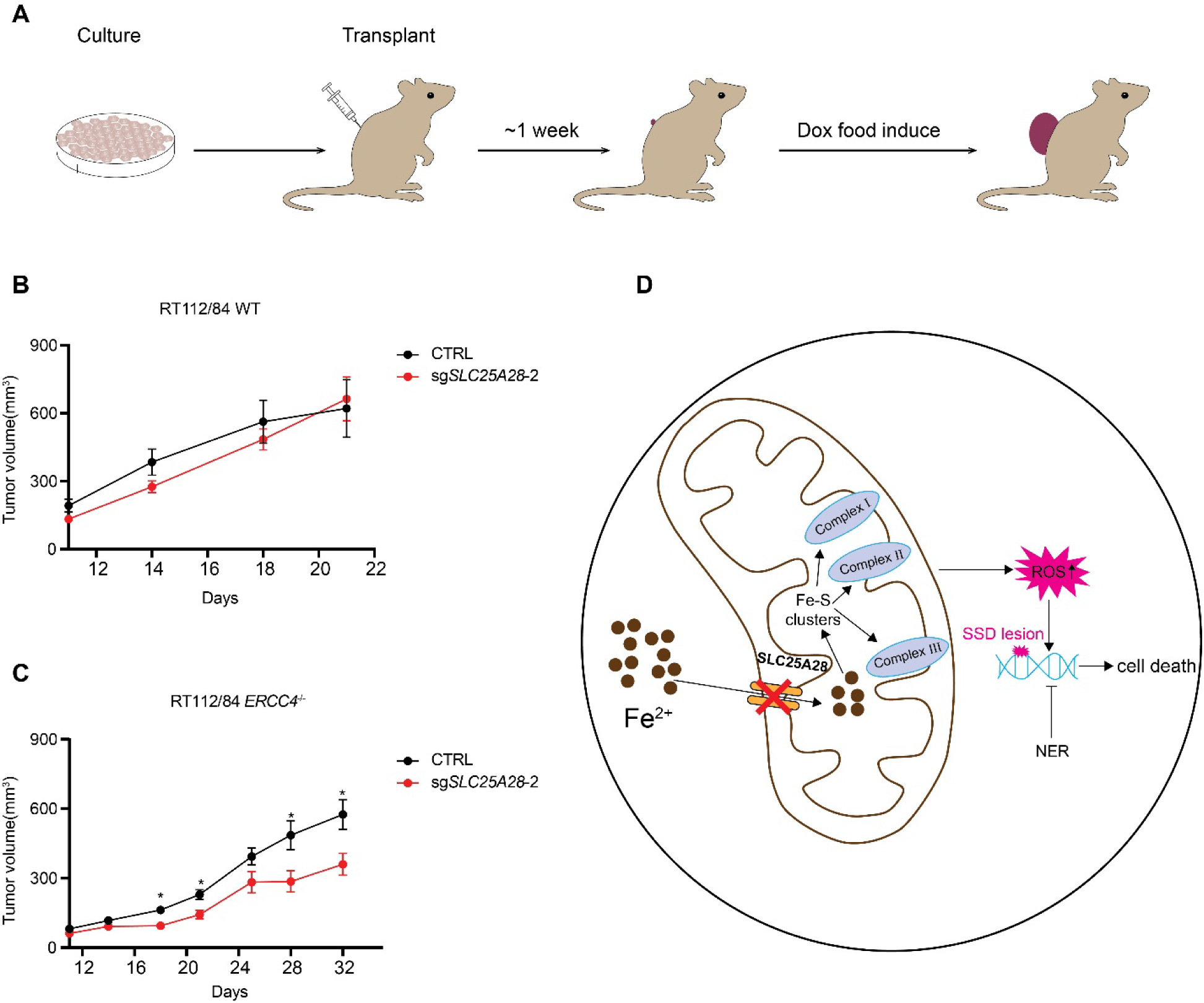
Synthetic-lethal interaction between *SLC25A28* and *ERCC4* in vivo and proposed mechanism. (A) Schematic of the xenograft experiments. (B–C) Tumor growth curves of xenografts generated from RT112/84 WT (B) or *ERCC4*-deficient (C) cells expressing sgNon-targeting (CTRL) or sg*SLC25A28*-2. Mice were switched to a doxycycline-containing diet to induce cas9 expression when tumors reached an average volume of ∼50–200 mm³. Data are presented as the mean ± SEM. (D) Proposed mechanism illustrating the interaction of *SLC25A28* with *ERCC2*, *ERCC3*, *ERCC4*, and *ERCC5*. *p<0.05.

Our data suggest that ROS accumulation is a central mediator of *SLC25A28* synthetic lethality. RNA-seq and GSEA analyses revealed enrichment of oxidative stress–related pathways, including hydrogen peroxide and general ROS response signatures, which were corroborated by DCFDA assays demonstrating increased intracellular ROS following *SLC25A28* knockout. These transcriptional and functional changes are consistent with elevated ROS associated DNA damage, as indicated by enrichment of Kyng DNA damage and UV response gene sets, potentially reflecting engagement of the NER pathway. Notably, DNA damage was not directly measured, and this arm of the model remains to be validated. Importantly, SLC25A28 encodes mitoferrin-2, a mitochondrial iron transporter required for iron-sulfur (Fe-S) cluster biosynthesis(45,46). While *SLC25A28* has been previously linked to vulnerabilities in chromosome 8p-deleted tumors due to paralog dependency with SLC25A37(32), our data reveal a novel context in NER deficiency. On the basis of these established functions, we propose a model in which loss of *SLC25A28* reduces mitochondrial iron import, leading to decreased Fe-S cluster biogenesis. Because Fe-S clusters serve as essential cofactors for multiple components of the mitochondrial electron transport chain(47), their depletion expected to impair electron transport, resulting in mitochondrial dysfunction and further ROS accumulation. This mechanistic link connects mitochondrial iron handling to the DNA damage burden specifically in NER-deficient cells. We emphasize that the Fe-S and electron-transport-chain steps of this model were not directly assessed in the present study. Supporting the broader role of *SLC25A28* in mitochondrial metal homeostasis, recent independent work has implicated it in cuproptosis, a copper-dependent regulated cell death pathway in breast cancer cells(48). Although cuproptosis was not directly interrogated in our study, these findings reinforce the concept that *SLC25A28* is a pivotal regulator of mitochondrial redox balance. Together, our integrated model posits that NER deficiency combined with *SLC25A28* loss leads to ROS accumulation, DNA damage, and excessive stress on NER repair pathways, explaining the selective vulnerability of NER-deficient cells.

Targeting *SLC25A28* and its NER deficiency interaction represents a mechanistically rational, though currently unproven hypothesis, synthetic lethal strategy for NER-deficient (NER-D) cancers. Distinct from conventional DNA-damaging platinum chemotherapeutics, exploiting this intrinsic metabolic vulnerability could, in principle confer selective toxicity toward NER-deficient tumors, potentially offering an approach to circumvent the limitations of current platinum chemotherapy. However, direct pharmacological targeting of SLC25A28 faces a fundamental obstacle: it is an inner mitochondrial multi-spanning membrane carrier with limited accessible pockets for small-molecule binding, posing substantial druggability challenges. Development of direct SLC25A28 inhibitors therefore remains aspirational and would require dedicated medicinal-chemistry effort. To circumvent this limitation, alternative therapeutic modalities warrant future exploration, such as pharmacological suppression of soluble enzymes responsible for mitochondrial iron–sulfur cluster biogenesis (e.g., NFS1 and ISCU) recapitulates mitochondrial iron deficiency(49), excessive ROS accumulation and the downstream synthetic lethal effect triggered by *SLC25A28* loss. Moreover, pharmacologically elevating intracellular oxidative stress through inhibiting glutathione biosynthesis or SOD2-mediated antioxidant defense selectively could conceivably exacerbate irreparable oxidative DNA damage in NER-deficient tumor cells while sparing NER-proficient normal tissues, thereby establishing a favorable therapeutic index. We emphasize, however, that these targets are broadly required for normal cellular and mitochondrial function; the assumption of a favorable therapeutic index and selective sparing of NER-proficient normal tissues is a hypothesis that remains to be tested and is not supported by data in the present study. Consistently, our genomic data encompassing 95,141 cancer patients supports inactivating mutations in *ERCC2/3/4/5* as a companion diagnostic biomarker for patient stratification, facilitating streamlined clinical trial design for therapeutics targeting iron metabolism or ROS homeostasis.

Our study has several limitations. The number of *ERCC4*-mutant cell lines in public datasets such as DepMap is limited, potentially constraining generalizability of the dependency analysis. *In vitro* models may not fully capture tumor heterogeneity, and xenograft experiments were conducted in immunodeficient mice, lacking immune context. Mechanistically, although we provide evidence linking *SLC25A28* loss to elevated ROS, DNA damage was inferred from transcriptional signatures rather than directly measured, and the proposed iron-sulfur cluster and electron-transport-chain intermediates were not assessed. additional biochemical validation could further establish causality.

Future work should validate *SLC25A28* synthetic lethality in additional NER-deficient tumor types and patient-derived xenograft models. Because SLC25A28 is a multi-spanning inner-mitochondrial carrier with limited small-molecule accessibility, direct inhibitor development may be challenging; pathway-level approaches targeting Fe-S biogenesis or redox homeostasis therefore warrant parallel exploration, together with assessment of pharmacodynamics and therapeutic window. Combination strategies with redox-modulating agents or DNA repair inhibitors may enhance efficacy. Finally, identification of predictive biomarkers for NER deficiency could enable patient stratification for targeted therapy.

In summary, our study identifies *SLC25A28* as a previously unrecognized, candidate synthetic lethal vulnerability in NER-deficient cancers, providing a rationale for precision therapeutic strategies that could circumvent the toxicity of platinum-based chemotherapy. More broadly, these findings highlight the potential of exploiting metabolic vulnerabilities in DNA repair–deficient tumors as a complementary therapeutic strategy.

## Authors’ Disclosures

No disclosures were reported.

## Authors’ Contributions

Nan Yang: Conceptualization, data curation, formal analysis, validation, investigation, visualization, methodology, writing–original draft, and writing–review and editing. Vijai Joseph: Conceptualization, analysis of patient data from the MSK-IMPACT database, writing, editing, illustrations and review of the manuscript. Lisa Hoeg and Daniel Durocher: Analysis of CRISPR screening data. Xuechun Bai and Jian Carrot-Zhang: Analysis of patient data from the MSK-IMPACT database. Kent W. Mouw: Provision of H460 WT and H460 *ERCC4^−/−^*cell lines. Kenneth Offit and Steven Lipkin: Supervision, funding acquisition, project administration, validation, and writing–review and editing.

## Acknowledgments

We would like to thank Xiaomu Wei and Junke Zhang for technical advice and support, and Zachary Nagel for providing the host cell reactivation assay plasmids as well as technical guidance. We also thank Jianmin Yang, Jesse Barnes, for their technical advice and support with the CRISPR screening experiments. We also acknowledge the following cores at MSKCC: Ralph Garippa of the Genome Editing and Screening Core, Elisa de Stanchina of the Antitumor Assessment Core, Nicholas D. Socci of the Bioinformatics Core, Ouathek Ouerfelli of the Organic Synthesis Core. This study was supported by STTR 1 R41 CA275627-01, the Niehaus Center for Inherited Cancer Genomics, and the Breast Cancer Research Foundation. We acknowledge the use of the Integrated Genomics Operation Core (RRID: SCR_027801), funded by the NCI Cancer Center Support Grant (CCSG, P30 CA008748).

## Notes

### Competing Interest Statement

The authors have declared no competing interest.

