## Supplementary Figures for "*SLC25A28* is a synthetic lethal vulnerability in nucleotide excision repair-deficient cancers"

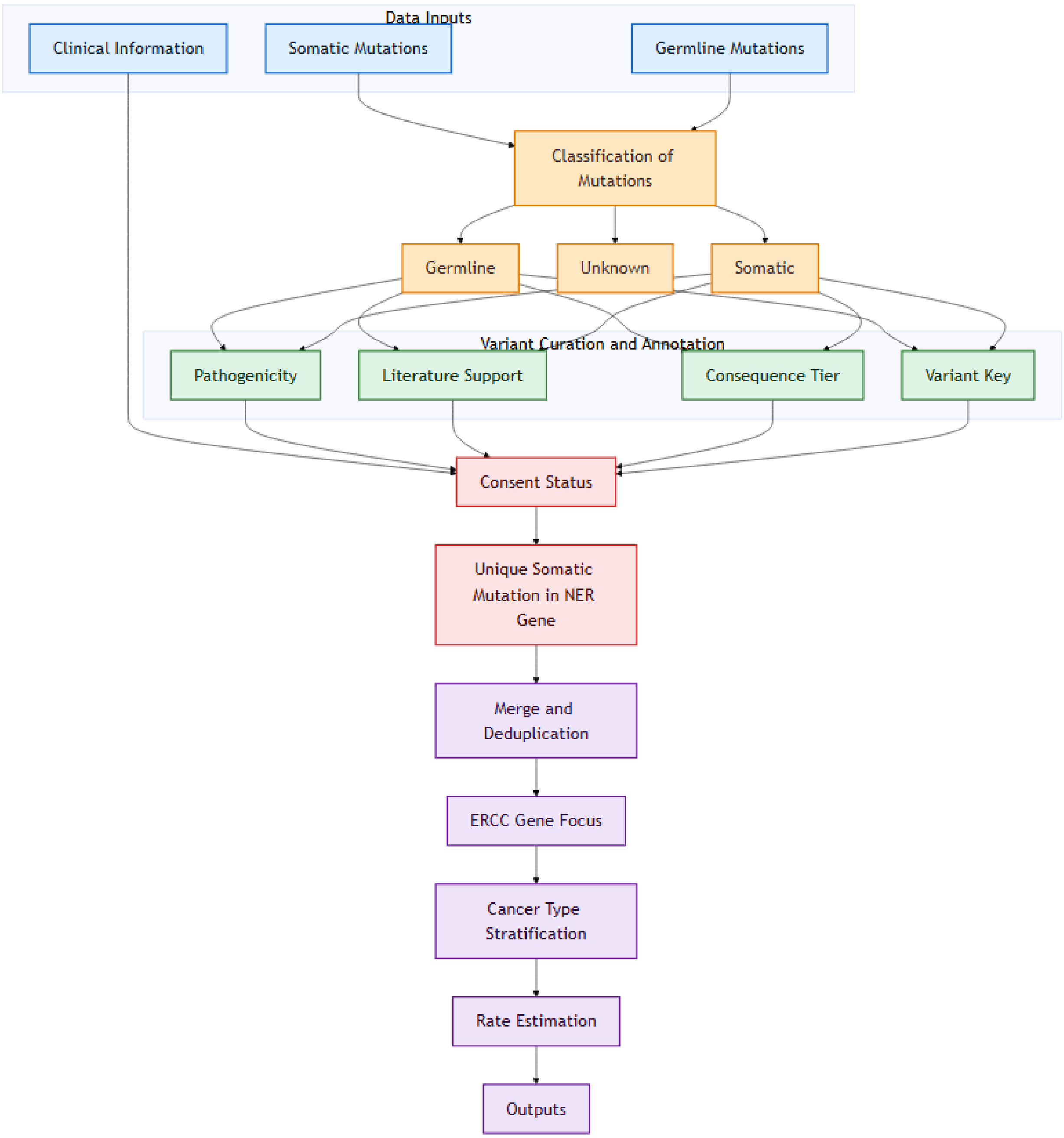


Figure S1. Variant filtering workflow for identifying patients carrying likely damaging *ERCC2*–*ERCC5* mutations, with alteration frequencies summarized across cancer types in a cohort of 95,141 patients with cancer profiled by MSK-IMPACT targeted sequencing.

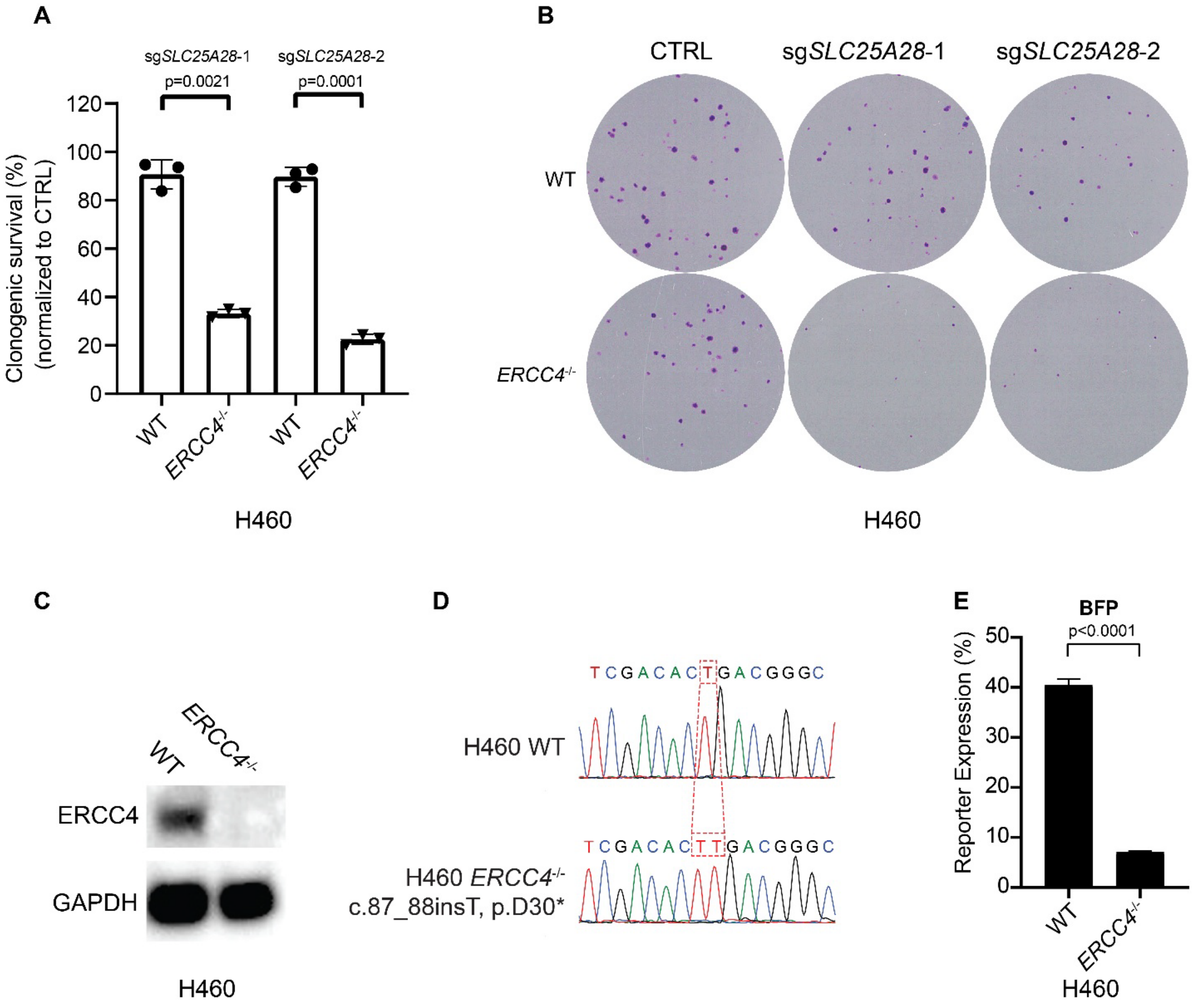


Figure S2. Identification of a synthetic-lethal interaction between *SLC25A28* and *ERCC4* in H460 cell lines. (A) Colony-formation assays of H460 WT and H460 *ERCC4*$^{-/-}$ cell lines transduced with the indicated sgRNAs or with sgNon-targeting (CTRL) lentivirus. Bars represent mean ± s.d. (n = 3). (B) Representative images of colony-formation assays corresponding to (A). (C) Western blot analysis shows loss of ERCC4 protein expression in H460 *ERCC4*$^{-/-}$ cells. (D) Sanger sequencing confirms the *ERCC4* frameshift mutation c.87_88insT (p.D30*) in the H460 *ERCC4*$^{-/-}$ cell line. (E) HCRA results demonstrate that the NER repair capacity of H460 *ERCC4*$^{-/-}$ cells is significantly reduced compared with that of H460 WT cells.

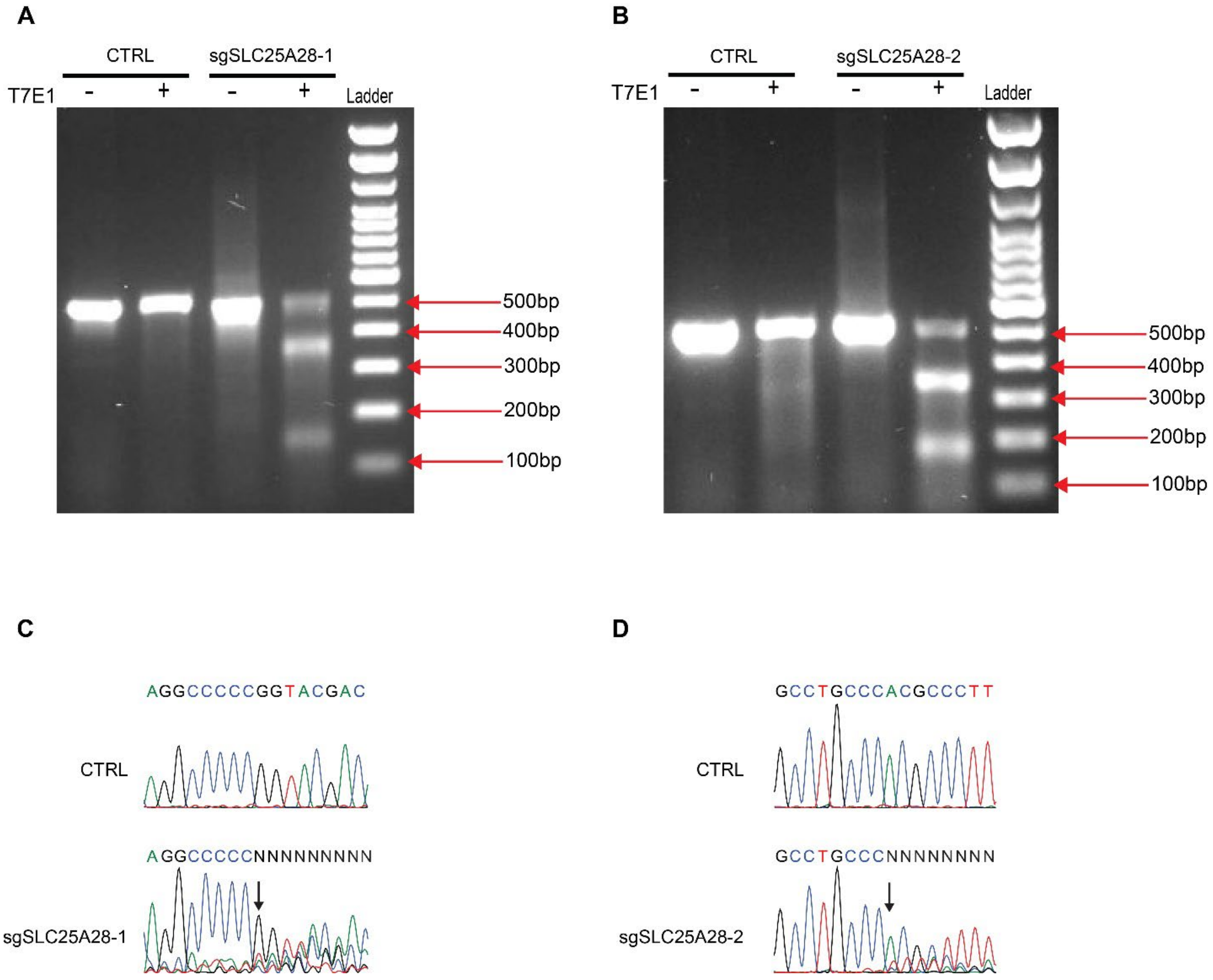


Figure S3. Validation of sgRNA-mediated editing efficiency of *SLC25A28*. (A, B) T7 endonuclease I (T7E1) assays of PCR amplicons spanning the *SLC25A28* target locus from Cas9-expressing RT112/84 WT cells transduced with sgNon-targeting (CTRL), sg*SLC25A28*-1 (A), or sg*SLC25A28*-2 (B). PCR products were treated (+) or not treated (−) with T7E1. (C, D) Sanger sequencing confirming indel formation at the *SLC25A28* locus in Cas9-expressing RT112/84 WT cells transduced with sg*SLC25A28*-1(C), or sg*SLC25A28*-2(D). Red arrows indicate the predicted Cas9 cleavage sites and the beginning of indel regions.

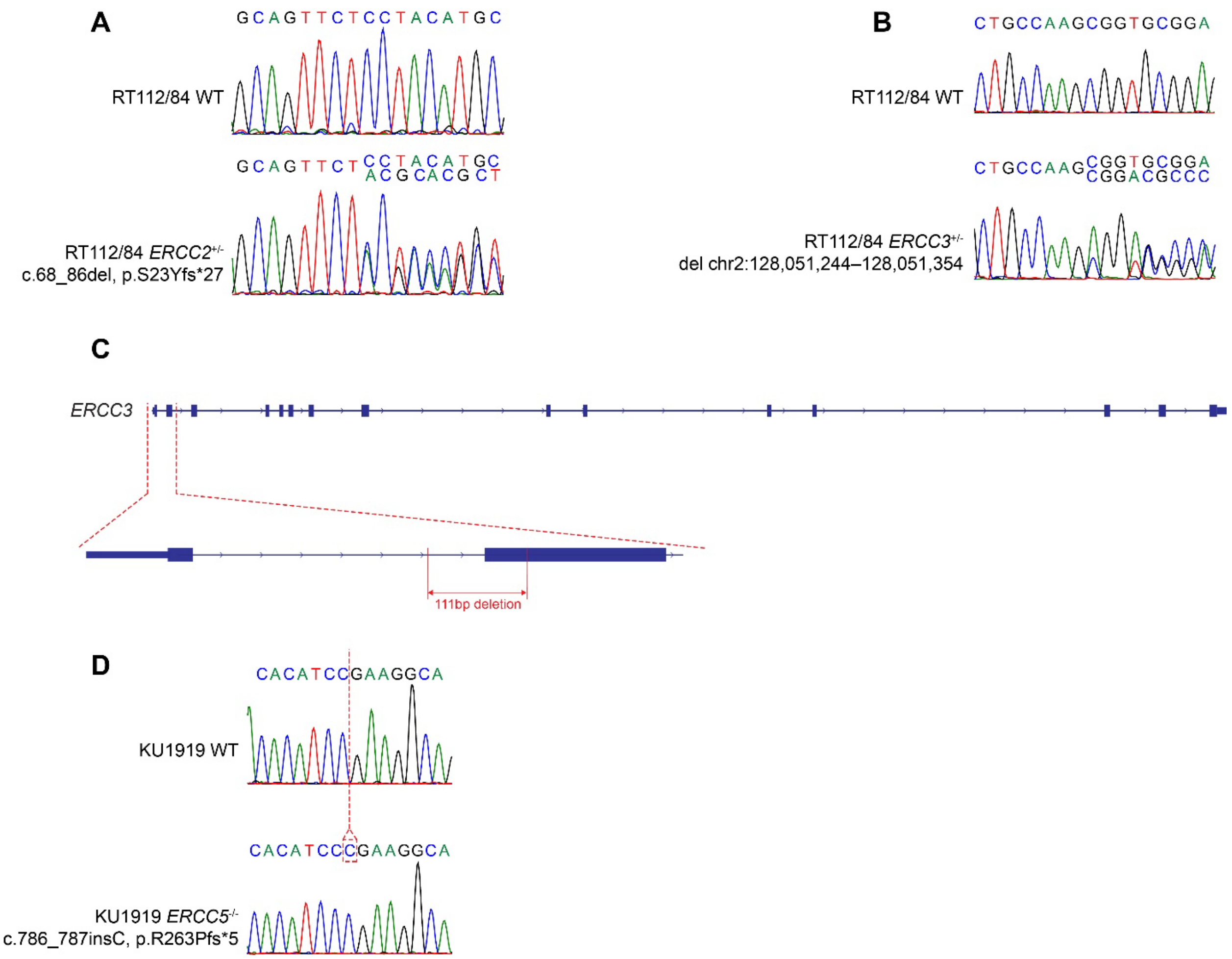


Figure S4. Sanger sequencing–based genotyping of RT112/84 *ERCC2*$^{+/-}$, RT112/84 *ERCC3*$^{+/-}$, and KU1919 *ERCC5*$^{-/-}$ cell lines. (A) Sanger sequencing confirms the *ERCC2* frameshift mutation c.68_86del (p.S23Yfs*27) in the RT112/84 *ERCC2*$^{+/-}$ cell line. (B) Sanger sequencing confirms the *ERCC3* deletion at chr2:128,051,244–128,051,354 in the RT112/84 *ERCC3*$^{+/-}$ cell line. (C) Schematic workflow illustrating a 111-bp deletion encompassing portions of intron 1 and exon 2 of *ERCC3* in the RT112/84 *ERCC3*$^{+/-}$ cell line corresponding to (B). (D) Sanger sequencing confirms the *ERCC5* frameshift mutation c.786_787insC (p.R263Pfs*5) in the KU1919 *ERCC5*$^{-/-}$ cell line.
